# Tegaserod maleate suppresses the growth of gastric cancer in vivo and in vitro by targeting MEK1/2

**DOI:** 10.1101/2022.04.01.486698

**Authors:** Zitong Wang, Yingying Chen, Xiaoyu Li, Yuhan Zhang, Xiaokun Zhao, Hao Zhou, Xuebo Lu, Lili Zhao, Qiang Yuan, Yunshu Shi, Jimin Zhao, Ziming Dong, Yanan Jiang, Kangdong Liu

## Abstract

Gastric cancer (GC), ranking fifth in global incidence and fourth in mortality. The current treatments for GC include surgery, chemotherapy, and radiotherapy. Although management and treatment strategies for GC have been improved over the last decade, the overall five-year survival rate remains less than 30%. Therefore, there is an urgent need to find novel therapeutic or preventive strategies that can increase GC patient survival rates. In the current study, we found that tegaserod maleate, a FDA-approved drug, could inhibit the proliferation of gastric cancer cells. Tegaserod maleate binds to MEK1 /2 and inhibits MEK1 /2 kinase activity. Moreover, the construction of CRISPER/Cas9 cell line further verified that tegaserod maleate depended on MEK1 /2 to inhibit the progress of gastric cancer. Notably, we found that tegaserod maleate suppressed tumor growth in patient-derived gastric xenograft (PDX) mouse model. We also compared tegaserod maleate with trametinib, a clinical MEK1/2 inhibitor, and comfirmed that tegaserod maleate have the same effect in inhibiting tumor volume and tumor weight. Our findings suggest that tegaserod maleate can inhibit GC proliferation by targeting MEK1/2.

## Introduction

Gastric cancer (GC) is characterized by insidious onset, asymptomatic or minor symptoms at an early stage, causing delayed diagnosis and poorer survival for most patients (1). At present, radical gastrectomy is the only potentially curative treatment for GC. Despite curative resection, recurrences occurrs in about 60% of patients. The main reason for this is that GC is usually advanced at the time of diagnosis. Preventing or reducing the frequency of the recurrence is probably more crucial than early detection of recurrence. (2, 3, 25). Hence, the development of novel drugs with low toxicity and high efficiency to prevent GC recurrence is a promising strategy to improve the survival rate of patients.

Patients with GC benefit from adjuvant therapy to reduce recurrence after curative resection. However, although postoperative chemoradiotherapy has an important effect on locoregional recurrence in patients with operable gastric cancer, there appears to be no additional benefit for regional and distant recurrence. Chemoprevention involves the use of natural or synthetic chemical agents to reverse, suppress or prevent the progression of a potentially malignant tumor to aggressive cancer and prevent cancer recurrence, thereby reducing cancer morbidity and mortality(4).For example, Celecoxib, a COX-2 inhibitor, has a significant effect on the regression of advanced gastric disease (6). Even with these encouraging results, the evidence for chemoprevention of gastric cancer recurrence are limited (5).

The development and approval of new drugs is a difficult and costly task with a high failure rate (8). Repurposing is attractive as a drug development strategy since the approved drugs are known including their drug similarity and pharmacokinetic characteristics, dosage, safety, and efficacy (9). Therefore, it is of great significance to screen the FDA approved drug library to select drugs with low toxicity and high efficiency to prevent cancer or recurrence. In recent years, it has been reported that tegaserod maleate, a 5-hydroxytryptamine 4 receptor (5-HT4R) agonist (10), can inhibit the proliferation of cancer cells (11,12). However, the inhibitory effect of tegaserod maleate on gastric cancer cells and its molecular mechanism remain unclear.

Here, we found that tegaserod maleate significantly suppressed the proliferation of GC in vitro and in vivo, and investigated the molecular mechanism of inhibition, providing reasonable theoretical support for the application of tegaserod maleate in the chemoprevention of gastric cancer.

## Materials and methods

### Reagents and antibodies

Tegaserod maleate (CAS:189188-57-6) was purchased from Selleck Chemicals LLC (Houston, TX, USA). Tegaserod maleate was dissolved with dimethyl sulfoxide to the final concentration was 50 mM and stored at -80°C. The following antibodies were used in the study: antiphospho-ERK1/2 (Thr202/Tyr204)(Cat# 4370, Cell Signaling Technology), Anti-ERK1/2 (Cat#4695, Cell Signaling Technology), anti-MEK1(Cat#2352, Cell Signaling Technology), anti-MEK2(Cat#9125, Cell Signaling Technology), anti-Ki-67 antibody (Cat#ab15580, Abcam), anti-Flag (Cat #F1804, Sigma).

### Cell culture

Human GC cell lines HGC27 and AGS were obtained from the Chinese Academy of Sciences (Beijing, China). HGC27 cell line was cultured in RPMI-1640 medium containing 10% fetal bovine serum at 37°C in a 5% CO_2_ humidified incubator. AGS cell line was maintained in F12K medium supplemented with 10% fetal bovine serum and cultured at 37°C in a 5% CO_2_ humidified incubator.

### Cell proliferation assay

GC cells (HGC27 cells: 2×10^3^ cells/well; AGS cells: 3×10^3^ cells/well) were seeded in 96-well plates. After incubation in the incubator for 16-18 h, the GC cells were treated with tegaserod maleate (0, 0.25, 0.5, 1, and 2 μM) for 24 h, 48 h, 72 h and 96 h. MTT (1mg/ml) was added at a ratio of 1:100. After incubation for 2 h, 100 μl DMSO was added terminate the reaction, and OD value at 490 nm/570 nm was measured with a microplate analyzer.

### Anchorage-independent cell growth assay

After the base layer agar was prepared with 0, 0.25, 0.5, 1 or 2 μM tegaserod maleate, HGC27 and AGS cells (8×10^3^ cells/well) were seeded in top layer agar with 0, 0.25, 0.5, 1 or 2 μM tegaserod maleate. The cells were cultured at 37°C in a 5% CO_2_ incubator for approximately 14 days. The colonies were counted by IN Cell Analyzer 6000 software.

### Anchorage-dependent cell growth assay

After seeding HGC27 (500 cells/well) and AGS (500 cells/well) in 6-well plates, cells were treated with 0, 0.25, 0.5, 1 or 2 μM tegaserod maleate. Two weeks later, the cells were fixed with 4% paraformaldehyde at 25°C for 30 min and stained with 0.1% crystal violet for 3 min.

### Western blotting

The HGC27 and AGS cells were inoculated into 10 cm dish. After 16 h, tegaserod maleate with different concentrations was added and treated cells for 24 h. The cells were harvested and lysed by Radio-Immunoprecipitation Assay (RIPA) buffer to obtain protein samples. Protein concentration were detected using the bicinchoninic acid protein assay kit (BCA Protein Assay Kit, Beyotime Biotechnology, Shanghai, China). Next, 30 μg protein extract was separated by sodium dodecyl sulfate-PAGE and transferred to polyvinylidene fluoride (PVDF) membranes. The membrane was blocked with 5% skim milk for 1h at room temperature. Next, the membranes were incubated overnight with the primary antibody at 4°C and then for 2 h with the secondary antibody at room temperature. Protein bands were displayed using the enhanced chemiluminescence (ECL) detection reagent (Dalian Meilun Biotechnology Co. Ltd, Dalian, China).

### Pull down assay

Epoxy-activated sepharose 4B conjugated with tegaserod maleate was prepared according to the manufacture’s instruction (#17-0480-01, GE Healthcare Life Sciences, Uppsala, Sweden). 200 μl tegaserod maleate-Sepharose 4B beads or vehicle were rotated with cell proteins (500 μg) in the reaction buffer. After rocking gently for 48 h at 4°C, the conjugated beads and vehicle were washed 4 times in the washing buffer followed by the addition of 30 μl 3× loading buffer at 100°C for 5 min. The binding was detected using Western blotting.

### In vitro kinase assay

Inactive ERK2 protein were incubated with active MEK1 recombinant protein (200 ng) or MEK2 recombinant protein (300 ng) for in vitro kinase assay. The reaction was run in kinase buffer (Cat#K02– 09, SignalChem, Canada) for 30 min at 30°C and stopped using 6 × loading. The protein was assessed using Western blotting.

### Cellular thermal shift assay

MEK1/2 was overexpressed in 293F cells. After 16 h, tegaserod maleate with was added and treated cells for 24 h. For the cell lysate CETSA, cultured cells were harvested and washed with PBS. The respective lysates were divided into smaller (100 μL) aliquots and heated respectively at different temperatures for 3 min followed by cooling at room temperature.for 3 min. The cell suspensions were frozen and thawed twice with liquid nitrogen. The lysate was separated from the cell debris by centrifugation at 12000 ×g for 20 min at 4°C. The supernatants were transferred to the new microtubes and analyzed by Western blotting

### Computer docking model

The Schrödinger Suite 2015 software program was used for silico docking. The three-dimensional (3D) structure of tegaserod maleate was derived from PubChem Compound. The Protein Preparation Wizard standard pipeline (Schrödinger Suite 2015) was used to process the structure of MEK1 (PBD:3EQG) and MEK2 (PBD:1S9I).

### CRISPR/Cas9 knockout cell lines

MEK1 /2 were deleted in GC cells using the CRISPR/Cas9 system. According to the manufacturer’s protocols, viral vectors and packaging vectors were transfected into 293T cells using Jet Primer (ThermoFisher Scientific, Waltham, MA, USA).. After 4 h, the medium was removed, and cells were cultured for 24 h, 48 h and 72 h. Viral particles were harvested and filtered using a 0.22 μm syringe filter, and then combined with 8 μg/mL polybrene to infect HGC27 and AGS cells. The cells were respectively treated with 2 μg/mL and 1 μg/mL puromycin for 72 h. Knockout efficacy was detected by Western blotting. The oligonucleotide sequences of MEK1 and MEK2 single guide (sg) RNA were listed as Table 1.

**Table 1.**
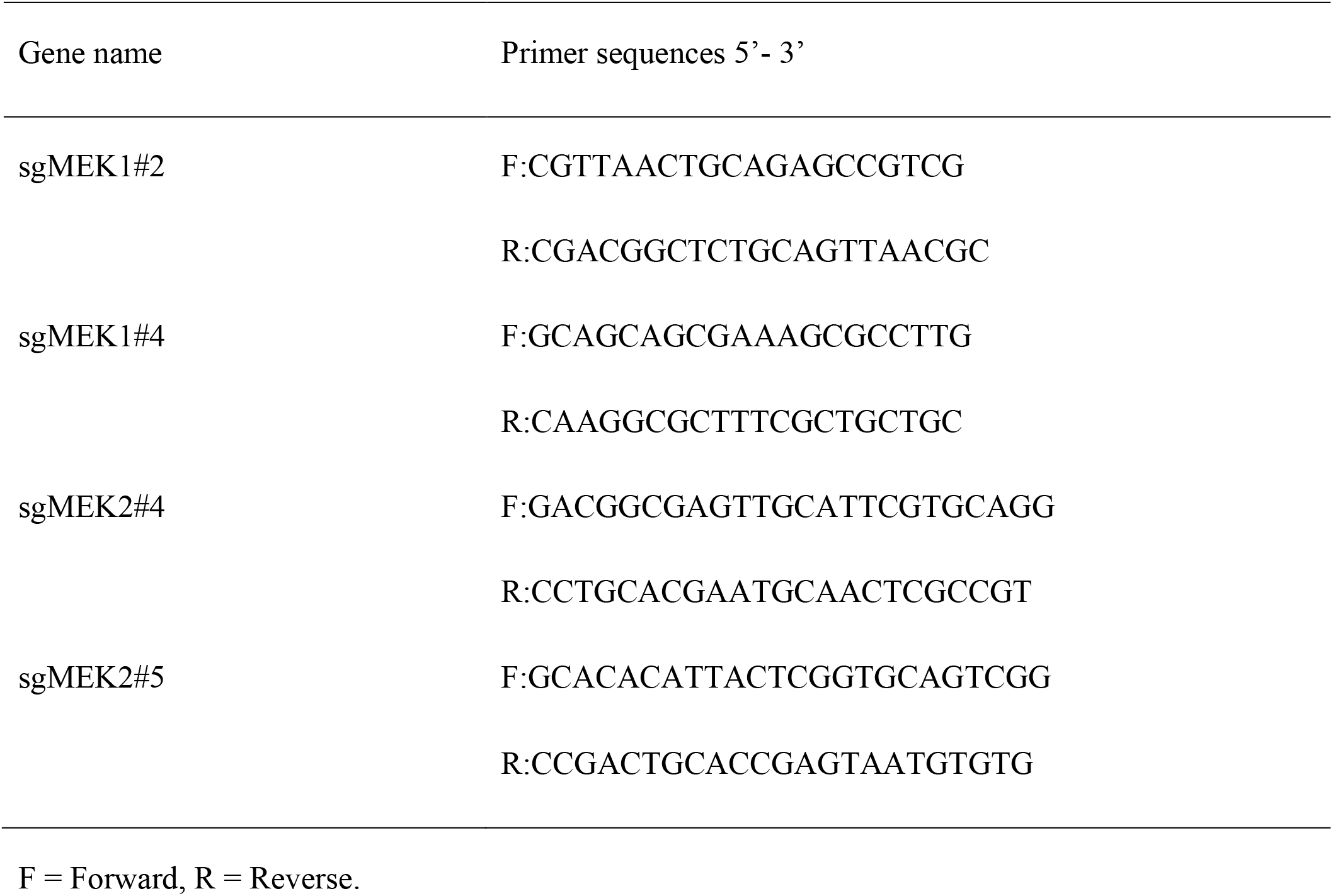
The oligonucleotide sequences of MEK1 and MEK2 single guide (sg) RNA.

### PDX mouse model

Severe combined immunodeficiency (SCID) mice were used (6 to 8 weeks old; Vital River Labs, Beijing, China) to investigate the effect of tegaserod maleate on GC PDX tumor growth, GC tissues were cut into about 1-2 mm pieces and planted into the back of the mice. The mouse were divided into three groups:vehicle group, low-dose group (2 mg/kg), high-dose group (10 mg/kg), with ten mice in each group. The vehicle or tegaserod maleate was orally taken once daily. Each mouse were checked twice weekly for the tumor volume and body weight. Until the tumor volume reached 1000 mm^3^, the tumor was extracted and then the tumor weight was measured.

### Immunohistochemistry (IHC) analysis

The tumor tissues were extracted and embedded in paraffin blocks for IHC staining. The tissue sections were deparaffinized, hydrated, antigen-repaired and blocked. Sections were incubated at 4°C with primary antibodies of Ki-67 overnight and then for 15 min with the secondary antibody at 37°C, following by diaminobenzidine and hematoxylin staining. Then, the sections were dehydrated and covered with slides. All tissue sections were taken with a microscope camera and analyzed using TissueFAXS Viewer software program.

### Statistical analysis

SPSS statistical software, version 21 (IBM Corp.) was used for all statistical analysis. The results were used non-parametric test or one-way analysis of variance (ANOVA) to compare significant difference. Quantitative data were expressed as mean values ± standard deviations. Statistical significance was defined by a p-value < 0.05.

## Results

### Tegaserod maleate inhibited GC cells proliferation *in vitro*

Tegaserod maleate, a FDA-approved drug, used for the treatment of irritable bowel syndrome(IBS-C). The chemical structure of tegaserod maleate is shown in **Figure 1A**. We first determined the toxicity of tegaserod maleate in GC cells by MTT assay, and found that the half-maximal inhibitory concentration (IC50) values of HGC27 cells at 48 h was 1.40 μM, and that of AGS was 2.14 μM (**Figure 1B**). Hereafter, we selected 0.25, 0.5, 1 and 2 μM of tegaserod maleate for further studies. Results revealed that tegaserod maleate (2 μM) inhibited HGC27 cell proliferation by 31.93 and 38.38 % at 72 and 96 h respectively, inhibited AGS cell proliferation of by 36.29 and 40.69 % at 72 and 96 h, respectively **(Figure 1C**). Then, we confirmed the anticancer effects of tegaserod maleate using anchorage-independent and anchorage-dependent cell growth assay **(Figure 1D&E**). Compared to the control group, results showed that the colony numbers of HGC27 and AGS cells were both decreased in a dose-dependent manner after tegaserod maleate treatment.. Hence, these results confirmed that tegaserod maleate could inhibit the GC cells proliferation *in vitro*.

**Figure 1.**
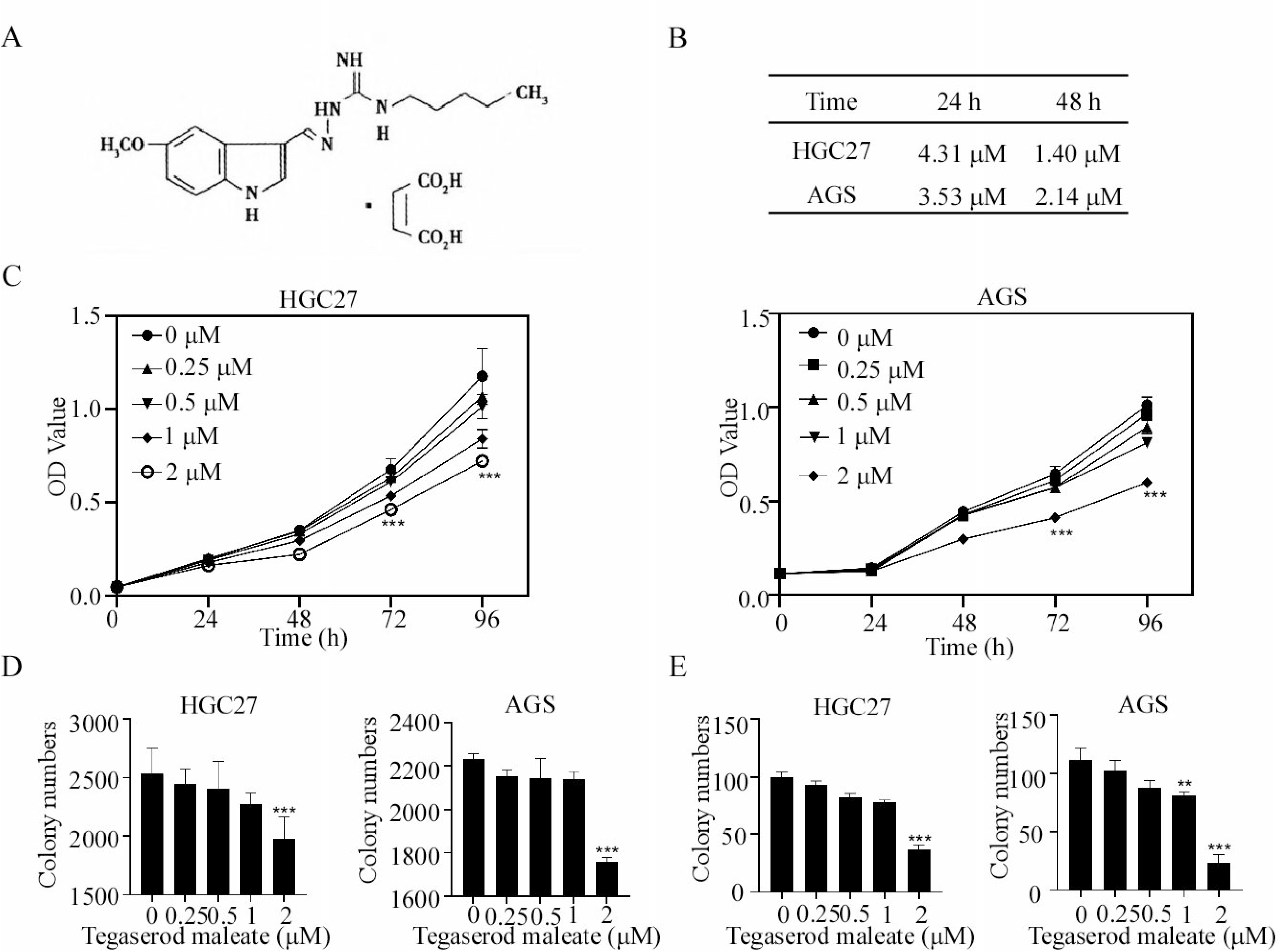
Tegaserod maleate inhibited GC cells proliferation *in vitro*. (A) Chemical structure of Tegaserod maleate. (B) Cell toxicity Assay. AGS and HGC27 treated with tegaserod maleate (0, 3.125, 6.25, 12.5, 25, 50 μM) for 24 and 48 h were evaluated by MTT. (C) The tegaserod maleate has an inhibitory effect on gastric cancer cells. AGS and HGC27 treated with tegaserod maleate (0, 0.25, 0.5, 1, 2 μM) for 0, 24, 48, 72 and 96 h were evaluated by MTT. (D) Tegaserod maleate inhibited anchorage-independent gastric cancer cells growth. AGS and HGC27 cells treated with tegaserod maleate (0, 0.25, 0.5, 1, 2 μM) for 2 weeks. (E) Tegaserod maleate inhibited anchorage-dependent gastric cancer cells growth. AGS and HGC27 cells treated with tegaserod maleate (0, 0.25, 0.5, 1, 2 μM) for 2 weeks. The asterisks (*) (**) (***) indicate a significant (P ˂ 0.05, 0.01 and 0.001).

### Tegaserod maleate could bind to MEK1 and MEK2

Since tegaserod maleate inhibited GC cells proliferation, we used in-silico docking assay to screen target. Our results indicated that tegaserod maleate could bind to MEK1 and MEK2. The results predicted that tegaserod maleate achieves binding with MEK1 at ASP208, MET146 and ASP152 sites, and with MEK2 at GLY83 and GLY81 sites (**Figure 2A**). Then, the computational docking results of tegaserod maleate with MEK1 and MEK2 were verified by pull down assay. We firstly attested that tegaserod maleate could bind to recombinant MEK1 and MEK2 protein ex vitro (**Figure 2B**). Then, we overexpressed MEK1 and MEK2 at HEK293F cell and found that Sepharose 4B-coupled-tegaserod maleate could also bind to MEK1 and MEK2 protein (**Figure 2C**). We then tested whether tegaserod maleate could bind to MEK1 and MEK2 in HGC27 lysate or AGS lysate. Results exhibited that Sepharose 4B-coupled-tegaserod maleate could bind to endogenous MEK1 and MEK2 protein (**Figure 2D**). Similarly, cellular thermal shift assay (CETSA) showed that the Tm values of the control group were 51.1 and 52.6°C, respectively, while the Tm values of the tegaserod maleate treatment group were 60.3 and 57.4°C, respectively, indicating that the Tm values shifted to the right, which further verified that the protein MEK1 and MEK2 in intact cells was more stable after tegaserod maleate treatment (**Figure 2E**). We then examined whether docking sites ASP208, MET146 and ASP152 of MEK1 and GLY83 and GLY81 of MEK2 <u>a</u>ctually make a difference. After the amino sites mutation, the cell lysate were harvested for pull down assay, and the resuls showed that the binding ability of tegaserod maleate was decreased(**Figure 2F**).These results suggested that tegaserod maleate bound to MEK1 /2.

**Figure 2.**
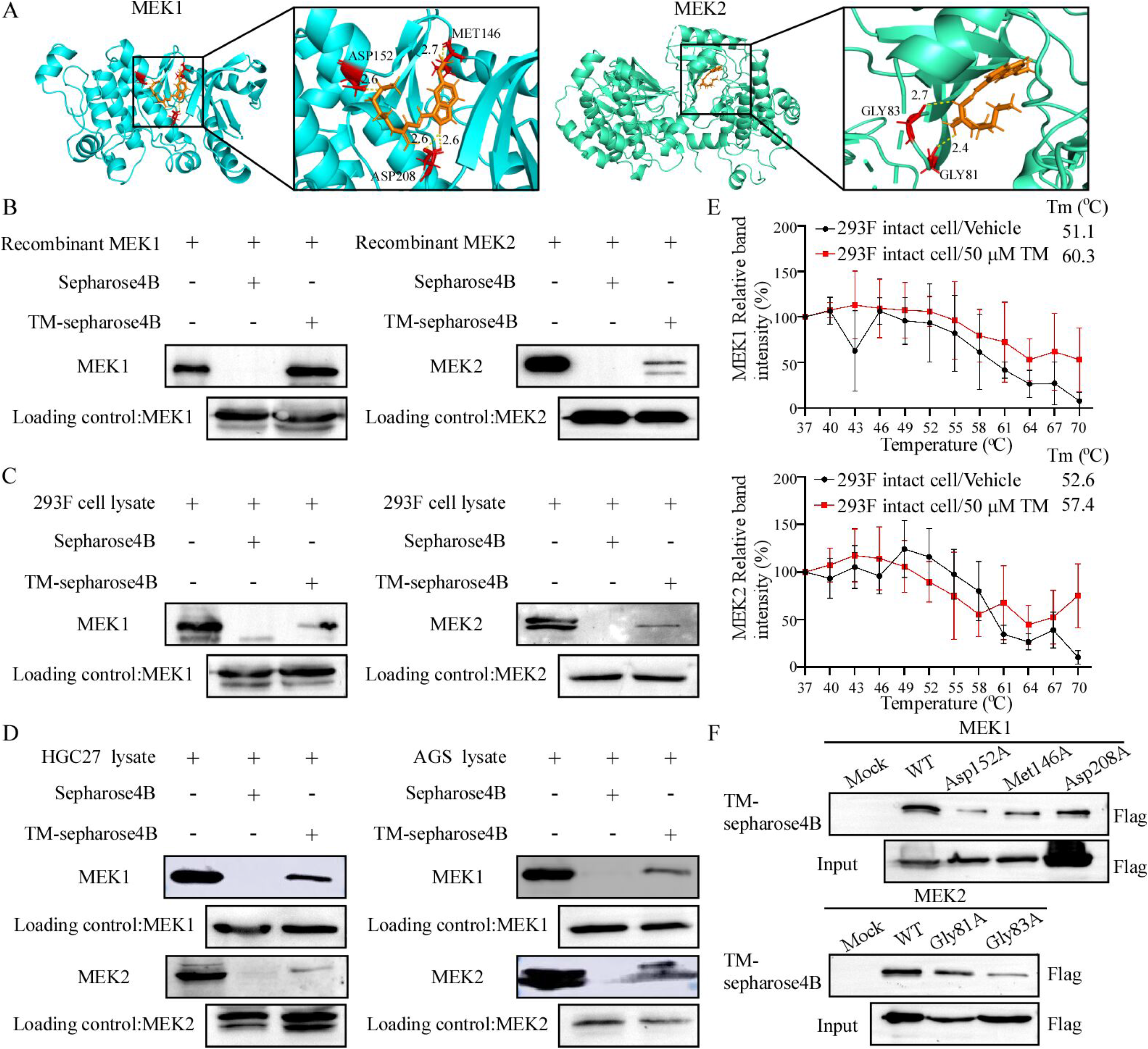
Tegaserod maleate could bind to MEK1/2. (A) Computational docking model between tegaserod maleate and MEK1 or MEK2. (B) The binding ability of tegaserod maleate to recombinant MEK1 and MEK2 protein, 293F cell which overexpressed MEK1 and MEK2 protein (C) and endogenic MEK1 and MEK2 protein in vitro (D), obtained via pull down assay. (E) Cellular thermal shift assay. The protein stability of MEK1 and MEK2 in intact cells. (F) Tegaserod maleate binding ability with mutant MEK1 and MEK2.

### Tegaserod maleate inhibited the MEK1/2-ERK1/2 signaling pathway in GC

Our above data indicates that tegaserod maleate could bind with MEK1 and MEK2. In vitro kinase assay was immediately performed. we found the phosphorylation efficiency of active MEK1 and MEK2 against inactive ERK2 was decreased at 0.25, 0.5, 1, and 2 μM tegaserod maleate **(Figure 3A)**. These results suggested tegaserod maleate could cause a decrease in MEK1/2 kinase activity and directly leaded to suppress ERK2 activation. Furthermore, we evaluated the downstream signaling molecule of MEK1 and MEK2. Results indicated that tegaserod maleate suppressed the phosphorylation of ERK1/2 in HGC27 and AGS cells. RSK2 is a key signaling molecule involved in cell proliferation and cancer development (13). After tegaserod maleate treatment, the downstream molecule RSK2 of ERK decreased in a dose-dependent pattern**(Figure 3B)**. Next, in order to verify that tegaserod maleate inhibited the activation of ERK by MEK1 and MEK2 in cell, we performed immunoprecipitate assay. Results showed that MEK1 and MEK2 could combine with ERK1/2 and tegaserod maleate could decreased the level of p-ERK1/2 T202/Y204 through suppressing the MEK1 and MEK2 kinase activity**(Figure 3C)**. In addition, the immunofluorescence assay was performed to investigate the variation of ERK1/2 T202/Y204 phosphorylation. The results showed that tegaserod maleate inhibited the fluorescence intensity of p-ERK1/2 T202/Y204 in a dose-dependent manner **(Figure 3D)**. Therefore, the above data supported that tegaserod maleate suppressed GC by blocking the MEK1/2-ERK1/2 pathway.

**Figure 3.**
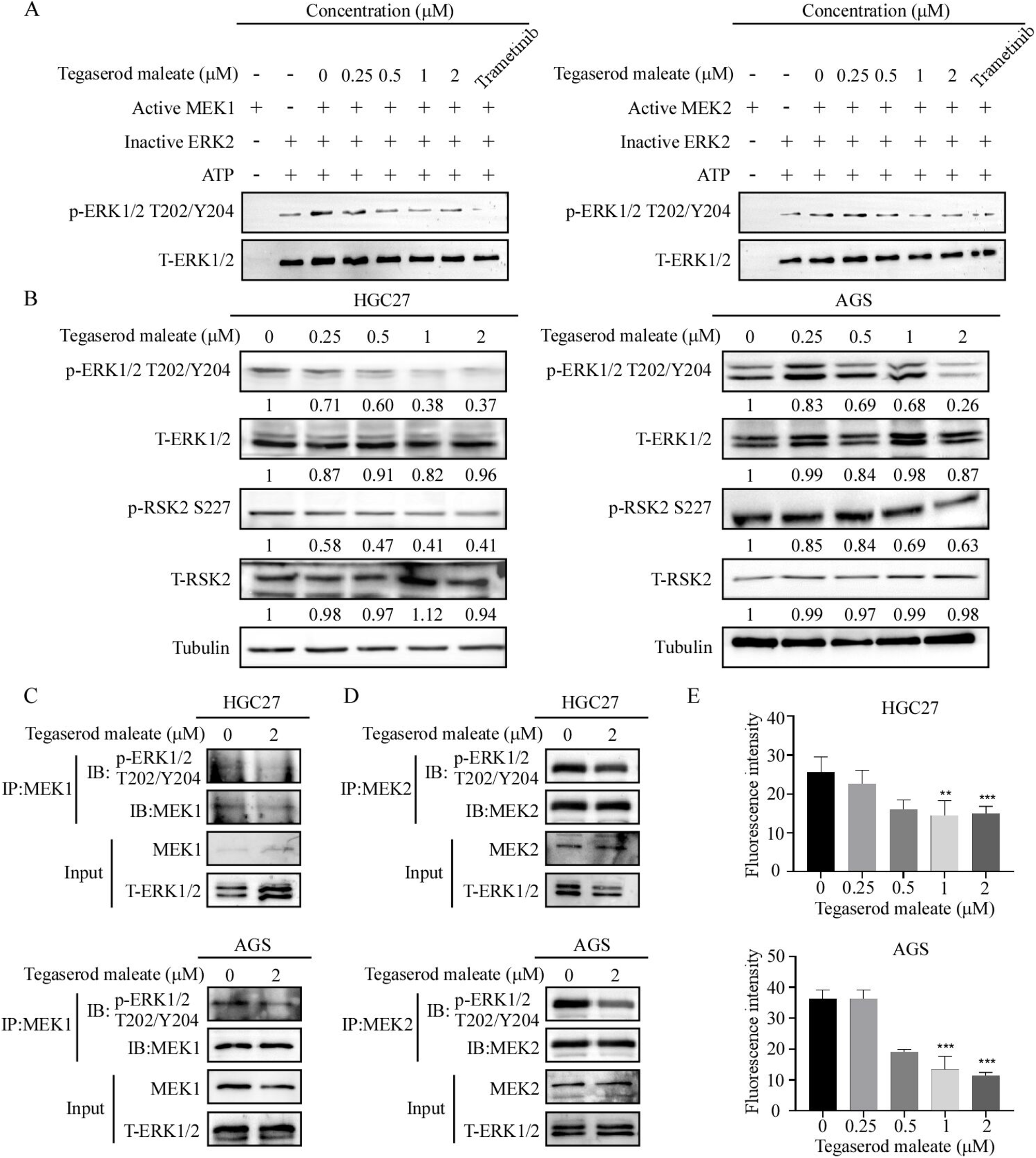
Tegaserod maleate inhibited the MEK1/2-ERK1/2 signaling pathway in GC. (A) MEK1 and MEK2 kinase activity was assessed by in vitro kinase assay using active MEK1, MEK2 and inactive ERK2 proteins. The effect of tegaserod maleate was determined by Western blotting analysis using a p-ERK1/2^T202/Y204^ antibody. (B) The level of p-ERK1/2, ERK1/2, p-RSK2 and T-RSK2 in HGC27 and AGS cells with different concentration of tegaserod maleate (0, 0.25, 0.5, 1 and 2 μM) treatment for 24 h was determined by Western blotting. The level of p-ERK1/2^T202/Y204^ was affected by MEK1 and MEK2 in HGC27 (C) and AGS (D) which treated with tegaserod maleate. ERK1/2 was immunoprecipitated by MEK1/2 and ERK1/2 was detected by p-ERK1/2^T202/Y204^. (E) Immunofluorescence staining of HGC27 and AGS: cells were treated for 24 h, and then stained for p-ERK1/2^T202/Y204^ (100 magnification).

### Tegaserod maleate inhibited gastric cell growth dependent on MEK1/2

To evaluate the role of MEK1 and MEK2 in GC progression, then AGS and HGC27 cell lines were used for knockout assays. Results indicated that sgMEK1#2, sgMEK1#4 reduced the protein levels of MEK1 significantly in both HGC27 and AGS gastric cell lines, while sgMEK2#4 and sgMEK2#5 could reduce MEK2 protein level (F**igure 4A)**. After knocking out of MEK1 and MEK2, the proliferation of HGC27 and AGS were suppressed **(Figure 4B)**. Similarly, the anchor dependent cell growth was also decreased in the knockout groups **(Figure 4C)**. Our data showed that tegaserod maleate could effectively inhibit the GC cells proliferation and colony formation. However, it is unclear whether the inhibitory effect of tegaserod maleate depend on levels of MEK1 and MEK2. Therefore, we treated MEK1 and MEK2 knockout GC cells with tegaserod maleate. After tegaserod maleate treatment for 96 h, cell viability was evaluated in HGC27 and AGS cell lines in mock and knockout cells groups to assess their sensitivity to tegaserod maleate. Results showed that the GC cells viability with tegaserod maleate treatment in the mock group was lower than those of sgMEK1#2 and sgMEK1#4 groups. Similarly, compared with the mock group, the cell viability of sgMEK2#4 and sgMEK2#5 was increased after tegaserod maleate treatment **(Figure 4D)**. These results suggested that tegaserod maleated does exert inhibitory effect depend on MEK1 and MEK2.

**Figure 4.**
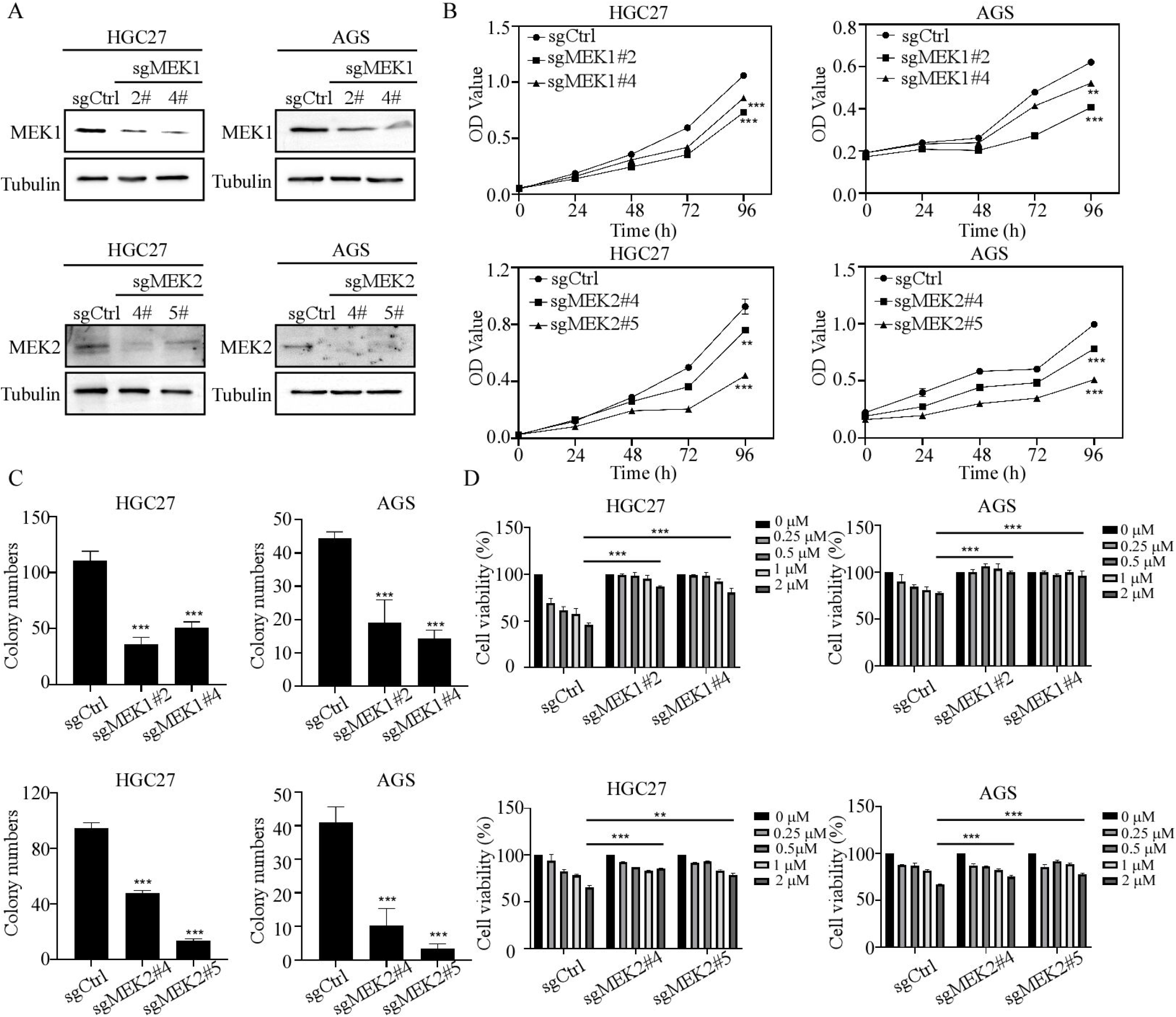
Tegaserod maleate inhibited gastric cell growth dependent on MEK1/2. (A) MEK1 and MEK2 knockdout efficiency was assessed in HGC27 and AGS cells. (B) Mock and sgMEK1 or sgMEK2 groups were deteted for 0, 24, 48, 72 and 96 h. Cell proliferation was evaluated by MTT assay. (C) Anchoring dependence ability of Mock and sgMEK1 or sgMEK2 groups. (D) Mock and sgMEK1 or sgMEK2 groups were treated with either tegaserod maleate or DMSO for 96 h. Cell viability was evaluated by MTT assay.

### Tegaserod maleate inhibited the growth of GC PDX *in vivo*

In order to evaluate the antitumor activity of tegaserod maleate *in vivo*, PDX mouse model of LSG85 and LSG51 cases with high MEK1 protein level were established from the GC PDX specimen repository**(Figure 5A&B)**. After tegaserod maleate was orally taken once daily, the tumor volumes of high-dose tegaserod maleate treatment group both in LSG85 and LSG51 cases were inhibited when compared with the vehicle group **(Figure 5C)**. After sacrificing the mice and removing tumors, the tumor picture in both LSG85 and LSG51 cases were shown in **Figure 5D**. We assessed tumor weight and detected that tumor growth inhibition (TGI) reached 34.32% in LSG85 case and 56.70% in LSG51 case with 10 mg/kg tegaserod maleate **(Figure 5E)**. In addition, tissue slice were prepared followed by Ki-67 staining for mechanism study. Immunohistochemical staining showed that the positive rates of the Ki-67 were reduced after tegaserod maleate treatment in LSG85 and LSG51 cases **(Figure 5F)**. Synchronously, we investigated the levels of p-ERK1/2 T202/Y204 in tumor tissues by Western blotting. These results showed compared with the vehicle group, these proteins were inhibited in the tegaserod maleate-treatment group (**Figure 5G**). In conclusion, tegaserod maleate suppressed GC tumor growth by inhibiting MEK1/2-ERK1/2 signaling pathway *in vivo*.

**Figure 5.**
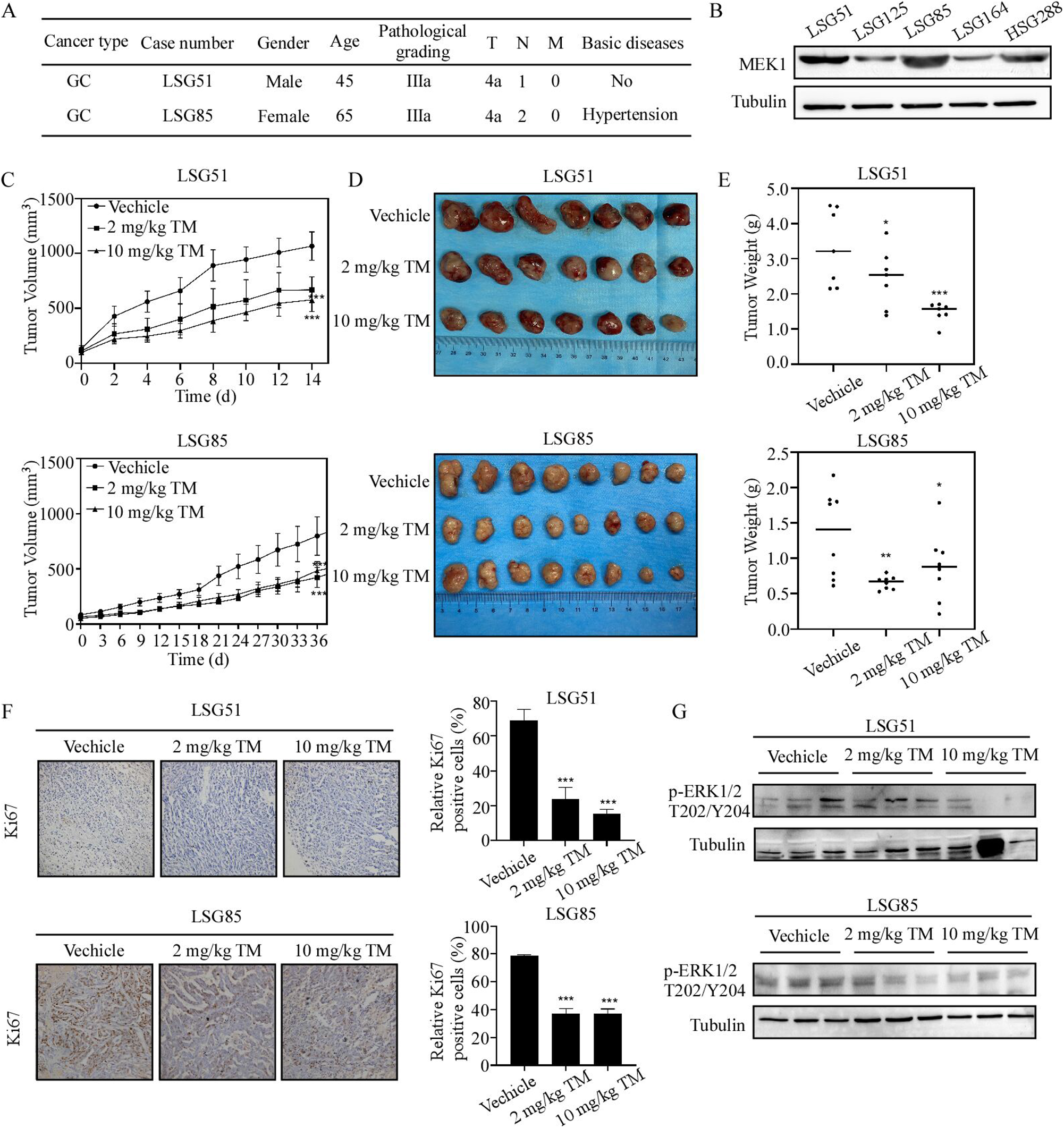
Tegaserod maleate inhibited the growth of GC PDX in vivo. (A) The information of two PDX cases. (B) The protein levels of MEK1 in different PDX cases. (C) The change of average tumor volume in different group of LSG85 and LSG51 cases after tegaserod maleate treatment. (D) Tumor images of different groups after sacrifice. LSG85 (n=8), LSG51 (n=7). (E) Tumor weight analysis in different groups of LSG85 and LSG51 cases after tegaserod maleate treatment compared with the average tumor weight of the vehicle group. (F) Left panel: Representative IHC images of LSG85 and LSG51 tumor tissue slices (100 magnifications), tumor tissues were stained with anti-Ki67; Right panel: Statistical analysis of IHC positive staining of Ki67 in both LSG85 and LSG51 cases. (G) The protein levels of p-ERK1/2^T202/Y204^ in tumor tissues were detected by Western blotting

### Tegaserod maleate have the same inhibitory effect compared with MEK1/2 inhibitor Trametinib

To confirm whether tegaserod maleate can be used as an inhibitor of MEK1 and MEK2 in gastric cancer. Here, we compared antitumor effects of tegaserod maleate with trametinib, a FDA-approved MEK1/2 inhibitor. The dosages of trametinib were selected based on previous studies. We selected HSG288 human gastric cancer tissues to established GC PDX model **(Figure 6A)**. The tumor volumes of both tegaserod maleate treatment groups and trametinib group in HSG288 case were also decreased **(Figure 6B)**. As shown in **Figure 6C**, tumor images of different groups after sacrifice. Subsequently, we assessed tumor weight and detected that TGI reached 68.41% with 10 mg/kg tegaserod maleate. Importantly, there was statistically insignificant difference between tegaserod maleate-treated group trametinib-treated group (**Figure 6D**). Furthermore, we examined the effect of tegaserod maleate on Ki-67 expression using IHC. The quantified IHC results were more likely to exhibit that the tegaserod maleate-treated groups decreased protein levels of Ki-67 **(Figure 6E)**. Meanwhile, the expression level of P-ERK1/2 T202/Y204 in tumor tissues was detected by Western blotting. The results showed that both tegaserod maleate-treated group trametinib-treated group could inhibit the expression level of P-ERK1/2 T202/Y204 **(Figure 6F)**. In conclusion, tegaserod maleate has the potential to be used in GC chemoprevention as a MEK1/2 inhibitor.

**Figure 6.**
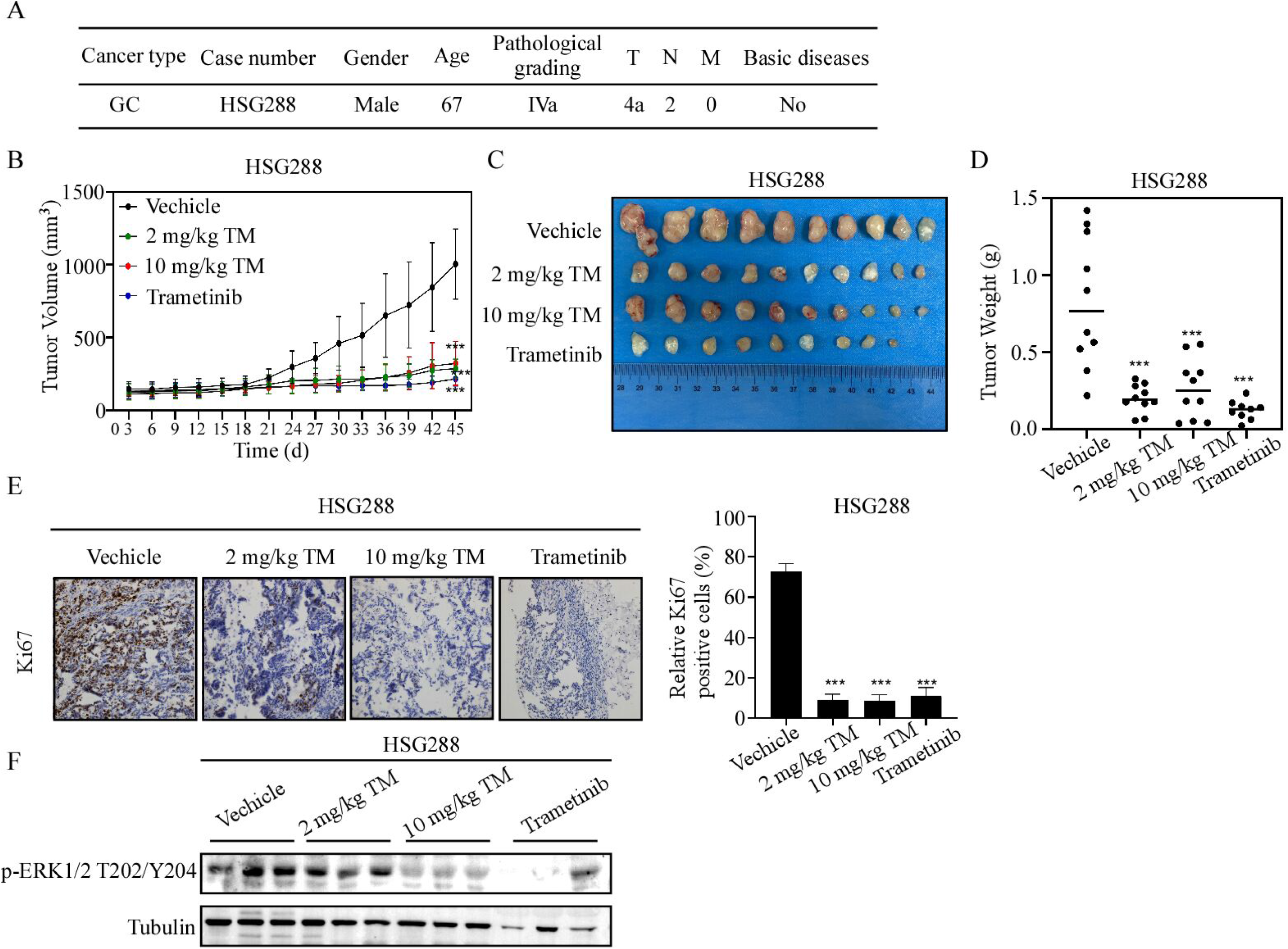
Tegaserod maleate have the same inhibitory effect compared with MEK1/2 inhibitor Trametinib. (A) The information of PDX cases. (B) The change of average tumor volume in different group of HSG288 case after tegaserod maleate and trametinib treatment. (C) Tumor images of different groups after sacrifice. HSG288 (n=10). (D) Tumor weight analysis in different groups of HSG288 case after tegaserod maleate and trametinib treatment. (E) Left panel: Representative IHC images of HSG288 tumor tissue slices (100 magnifications), tumor tissues were stained with anti-Ki67; Right panel: Statistical analysis of IHC positive staining of Ki67 in HSG288 case. (F) The protein levels of p-ERK1/2^T202/Y204^ in tumor tissues were detected by Western blotting.

## Discussion

Despite improvements in adjuvant therapies and general progress in oncogenesis mechanism comprehension for gastric cancer in recent years, the long-term survival of gastric cancer patients is not substantially improved due to high recurrence rates and a dismal prognosis(14, 26). As a result, chemoprevention strategies still appear as an effective strategy in reducing the incidence and the mortality for GC cancer. In recent years, there has increasing evidence demonstrates that drugs targeting nutrient metabolism are promising candidates for preventing tumourgenesis and cancer recurrence because of their availability and relatively low side effects compared to other chemotherapeutic drugs (21). Notably, many FDA-approved drugs that target nutrient metabolism, such as metformin, thiazolidinediones, statins, specific amino acid deprivation drugs and drugs targeting protein degradation have shown significant antitumor effects in various types of cancer (15). Moreover, by screening FDA approved drug library, our previous study confirmed that tegaserod maleate suppresses the proliferation of ESCC by blocking peroxisome function and the features of tegaserod maleate make it promising candidates for cancer chemoprevention (22). However, the underlying molecular targets and related mechanism still need further illustration. In our study, we identified that tegaserod maleate also exhibited anticancer effects on GC cells. Moreover, we confirmed that tegaserod maleate had a obvious inhibitory effect on gastric cancer PDX model and there was no statistically significant difference in inhibiting tumor volume or tumor weight compared with trametinib. According to clinical studies, trametinib, FDA-approved MEK1/2 inhibitor, was used in tumors with BRAF-activating mutations, including melanoma, non-small cell carcinoma, and thyroid cancer. However, trametinib has not been used in the clinical treatment of gastric cancer yet (23). Consequently, our findings suggested that MEK1 and MEK2 may also be targets for the prevention of gastric cancer and supported tegaserod maleate could be used in GC chemoprevention as a MEK1/2 inhibitor.

The MAP kinase cascade is the most important oncogenic driver of human cancers, and blocking this signaling module by targeted inhibitors is a promising antitumor strategy (16). Since MEK1/2 have very narrow substrate specificity, MEK1/2 inhibition specifically shuts off ERK1/2 signaling without affecting other signaling pathways. Therefore, inhibition of MEK1/2 is an effective method to suppress the whole cascade reaction and an efficient target for anticancer therapy (17). Futhermore, we detected the role of MEK1 and MEK2 in GC growth and found that the MEK1 and MEK2 protein levels were higher in GC (supplementary fig 2). In addition, the protein levels of MEK1 and MEK2 were negatively correlated with patient survival rate in GC (supplementary fig 2). Combined with MEK1 and MEK2 knockdout assays in GC cells, results identified that tegaserod maleate inhibited GC proliferation through MEK1 and MEK2. In present study, we utilized computational docking model to indicate that tegaserod maleate could bind to MEK1 /2 (Figure 2). Then, we also verified this by pull down assay and CETSA assay (Figure 2). The homology of peptide sequence of MEK1 (45 kDa) and MEK2 (46 kDa) encoded by MAP2K1 and MAP2K2 respectively was 85%, and the homology of catalytic domain was 86%. The typical MEK1/2 secondary structures consists of the N-terminal ∼70 amino acid residues, the protein kinase domain (∼290 amino acids), and the C-terminal ∼30 amino acid residues (20). The D208, M146 and D152 were located in the kinase domain of MEK1, and G81, G83 were located in the kinase domain of MEK2. Our results indicated that tegaserod maleate could bind with MEK1 at D208, M146 and D152 and MEK2 at G81, G83 respectively (Fig.2). In addition, utilizing kinase assay, we confirmed that tegaserod maleate suppressed the MEK1 and MEK2 kinase activity in vitro rather than detecting the signal only in cells. These data indicate that tegaserod maleate is an inhibitor of MEK1/2.

In conclusion, our finding attested that MEK1and MEK2 are promising therapeutic target for GC. Tegaserod maleate could bind to MEK1and MEK2 and inhibit its kinase activity. Tegaserod maleate also suppressed GC growth in vitro and in vivo by blocking MEK1/2/-ERK1/2 signaling pathway. Our study suggests that the proper application of tegaserod maleate may be a beneficial therapeutic strategy for GC patients with high MEK/2 levels.

## Acknowledgements

Thanks to Yifei Xie, Mingzhu Li, Xinyu He, Ang Li and Yin Yu from the China-US (Henan) Hormel Cancer Institute for their assistance.

## Author contributions

KL and YJ conceived the study and revised the paper. ZW performed most of the experiments and wrote the paper. YJ directed the experiment and proofread the manuscript. YC and XL analysed research data. YZ, XZ and QY contributed reagents or other essential material.YC, AL and LZ did immunohistochemistry. HZ and XL performed animal assays. All authors contributed to the article and approved the submitted version

## Data availability

The data supporting the findings of this study can be found in the article, Supplementary Information or available from the corresponding author upon reasonable request.

---

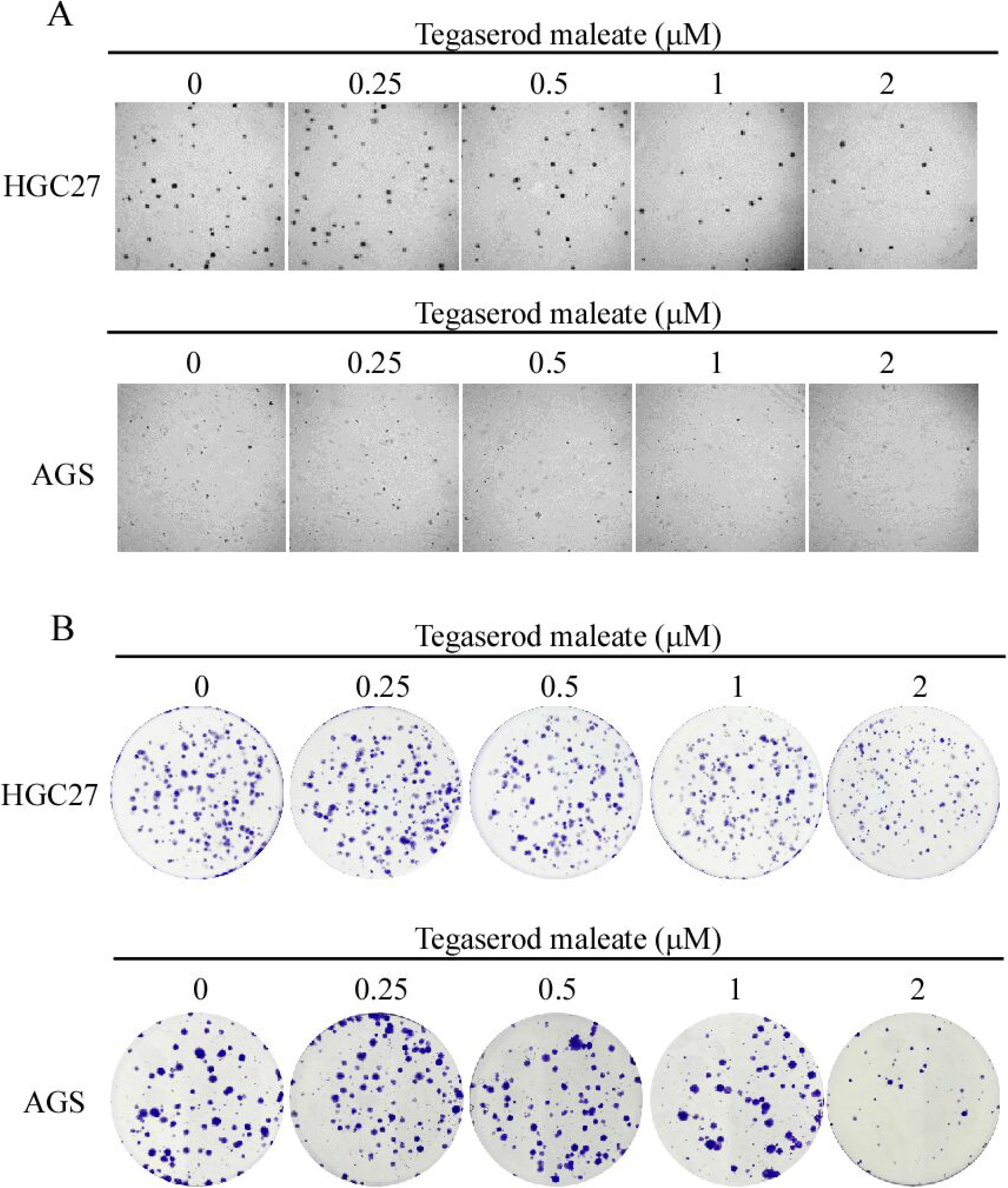

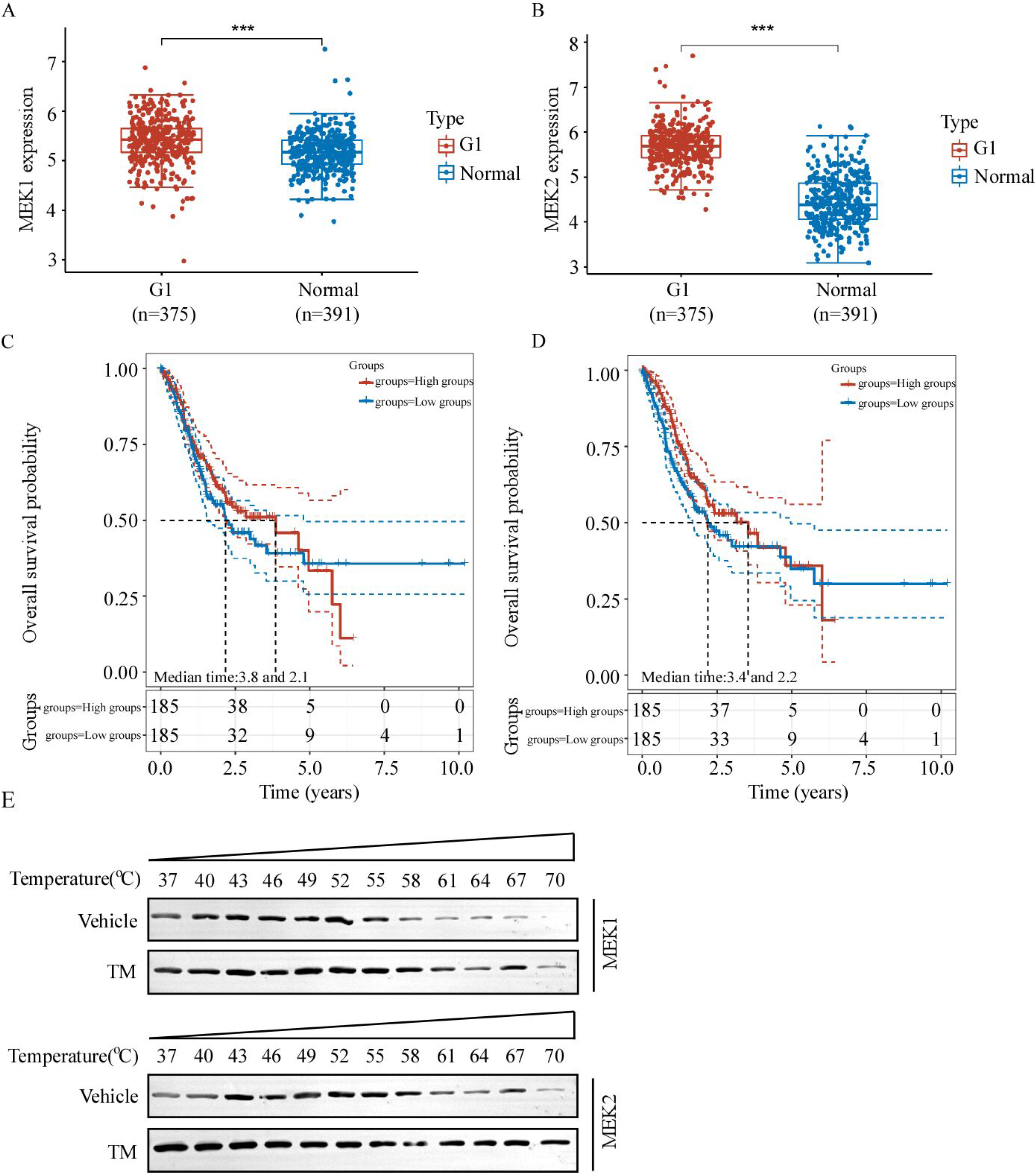

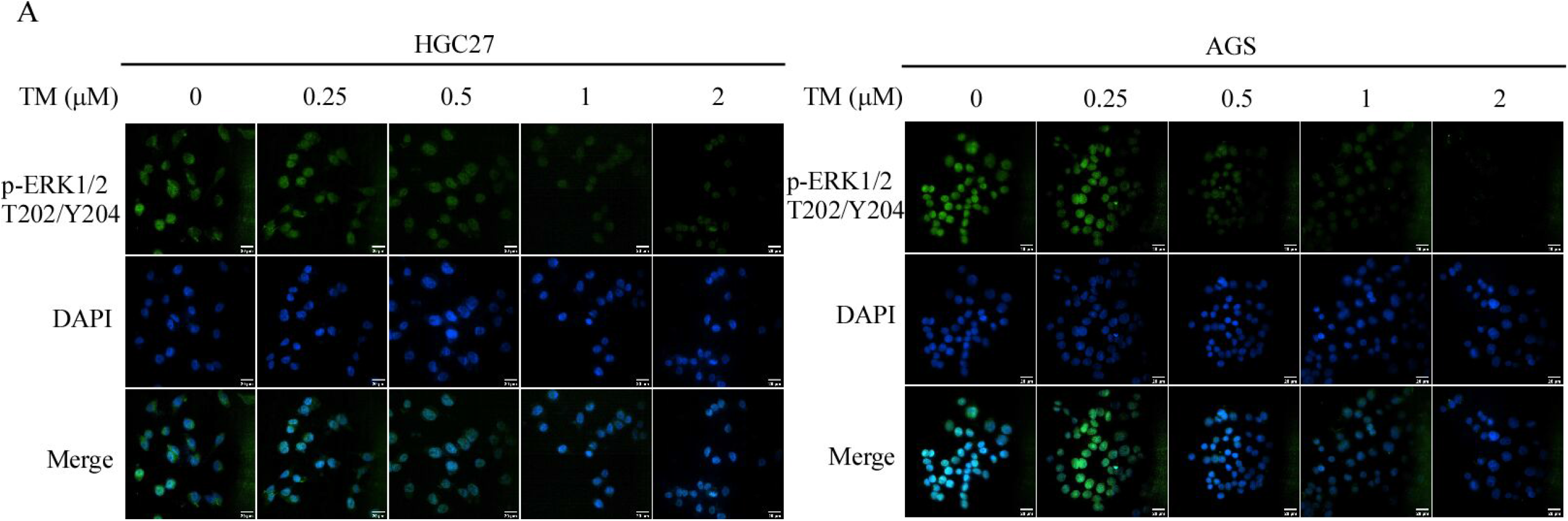

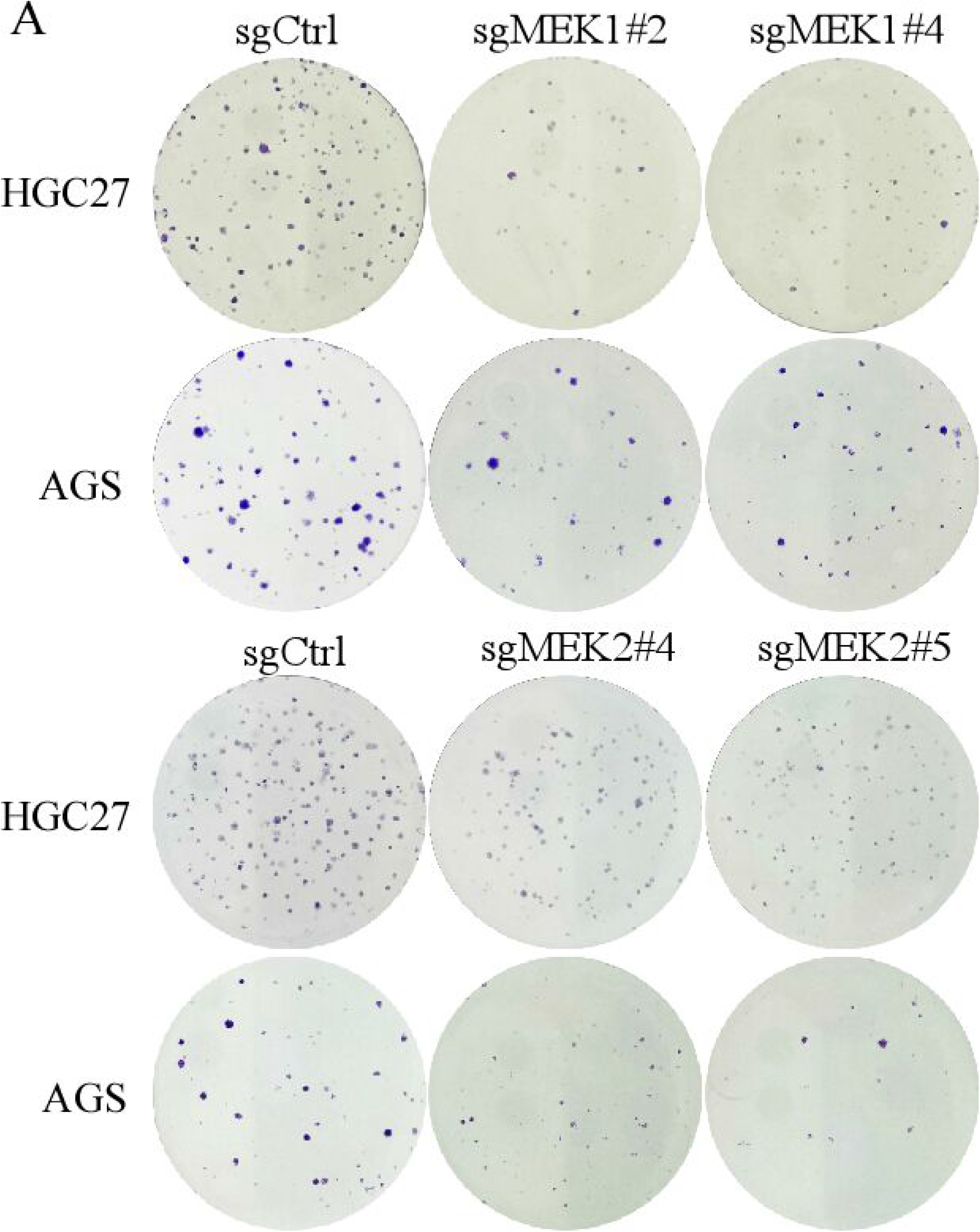

